# New tyrosinases with a broad spectrum of action against contaminants of emerging concern: Insights from *in silico* analyses

**DOI:** 10.1101/2020.07.30.225821

**Authors:** Marcus V. X. Senra, Ana Lúcia Fonseca

## Abstract

Tyrosinases (EC 1.14.18.1) are type-3 copper metalloenzymes with strong oxidative capacities and low allosteric selectivity to phenolic and non-phenolic aromatic compounds that have been used as biosensors and biocatalysts to mitigate the impacts of environmental contaminants over aquatic ecosystems. However, the widespread use of these polyphenol oxidases is limited by elevated production costs and restricted knowledge on their spectrum of action. Here, six tyrosinase homologs were identified and characterized from the genomes of 4 widespread freshwater ciliates using bioinformatics. Binding energies between 3D models of these homologs and ~1000 contaminants of emerging concern (CECs), including fine chemicals, pharmaceuticals, personal care products, illicit drugs, natural toxins, and pesticides were estimated through virtual screening, suggesting their spectrum of action and potential uses in environmental biotechnology might be considerably broader than previously thought. Moreover, considering that many ciliates, including those caring tyrosinase genes within their genomes are fast-growing unicellular microeukaryotes that can be efficiently culturable at large-scales under *in vitro* conditions, should be regarded as potential low-cost sources for the production of relevant biotechnological molecules.

## 1. Introduction

Contamination of aquatic environments with toxic compounds has become a major global threat and one of ours greatest challenges toward biodiversity and hydric recourses conservation (Galindo-Miranda et al., 2019). Contaminants of emerging concern (CECs) are pervasive and unregulated pollutants of natural and synthetic origin, which include fine chemicals, dyes, pharmaceuticals, cosmetics, personal care products, cyanotoxins, and pesticides among many others (Sauvé et al., 2014; Visanji et al., 2020). CECs usually reach aquatic compartments via domestic and industrial sewage or agriculture runoffs (Lapworth et al., 2012), where they are detected within nano (ng.L^−1^) or micro (µg.L^−1^) scales (Wilkinson et al., 2017) and frequently associated with a variety of short and long-term detrimental effects over biodiversity, community structure and human health (Brühl and Zaller, 2019; Lapworth et al., 2012; Saaristo et al., 2018; Sanchez and Egea, 2018; Santangeli et al., 2019). Because of the general inefficiently of conventional water treatment systems in the removal of such potutants (Bolong et al., 2009), the development of novel efficient and cost-effective technologies able to detect and/or remove the great diversity CECs from natural systems is greatly desirable (Bilal et al., 2019).

Tyrosinases (EC 1.14.18.1) are widespread polyphenol oxidases of bacteria, fungi, plants, and animals (Zaidi et al., 2014) that participate of critical biological functions, such as photoprotection (mammals), immune system and wound-healing processes (plants, fungi, and insects), and in spore and sexual organs maturation (fungi) (Agarwal et al., 2019). These metalloenzymes carry a characteristic type-3 copper center formed of two copper ions (CuA and CuB), each one connected to three conserved histidine residues (Goldfeder et al., 2014), which catalyzes hydroxylation monophenols to diphenols (monophenolase reaction) and oxidation diphenols to quinones (diphenolase reaction), using molecular oxygen as electron acceptor in both reactions (Kaintz et al., 2014). This process occurs as a catalytic cycle, where the active site shifts from active *oxy* (Cu^II^ - O_2_ - Cu^II^) state to intermediate *met* (Cu^II^ - OH - Cu^II^) state, and then, to a resting *deoxy* (Cu^I^ - Cu^I^) state that requires a new dioxygen for reactivation (Ba and Vinoth Kumar, 2017).

This biocatalyst have been largely used in food, cosmetic and pharmaceutical industries (Agarwal et al., 2019) and more recently, because of its low allosteric specificity and strong oxidative capacities over a diversity of phenolic and nonphenolic aromatic compounds (Asgher et al., 2014), emerged as a versatile biosensor and biocatalyst for monitoring and removing environmental contaminants, such as cresols, chlorophenols, phenylacetates, and bisphenols from natural systems (Ba and Vinoth Kumar, 2017; Harms et al., 2011). However, despite this great potential to mitigate the environmental impacts of CECs, most studies remain restricted to commercially available enzymes, purified from bacteria or fungi and to tests against a narrow diversity of pollutants. Moreover, their high production cost limits the widespread use of tyrosinases in field applications (Ba and Vinoth Kumar, 2017).

Bioinformatic tools represent a powerful, fast and cost-effective method for enzyme discovery (Vanacek et al., 2018). As a consequence of diffusion high-throughput sequencing technologies thousands of genomes from model and non-model organisms from all over the tree of life are now available within public sequence databases, which in turn have become a fruitful source of biological information. Despite of extensively used in drug discovery processes, including for detection and rational design of novel molecules and prediction of their potential cellular and sub-cellular targets (Xia, 2017), genomics tools (protein 3D structure modeling, molecular dynamics (MD), and docking assays) have only sparsely used in environmental biotechnology applications for identification and characterization of novel biocatalysts and prediction of potential targets. In an interesting work, Srinivasan and coworkers (Srinivasan et al., 2019) applied a molecular docking approach to indicate a potential a broad spectrum of action of laccases (EC 1.10.3.2) (polyphenol oxidases) against a diversity of textile dyes, and in a different study, Ahlawat and collaborators (Ahlawat et al., 2019), corroborated this data using MD and docking assays, but also validated their results through experimental procedures, highlighting potential applications of bioinformatics to speed up and reduce costs related to development of novel biocatalysts

Here, we screened the genomes of 23 ciliate species (Ciliophora) for the presence of tyrosinase homologs. Ciliates are ancient (~1180 Ma) (Fernandes and Schrago, 2019), highly diversified (> 8,000 valid species), and widespread unicellular microeukaryotes able to colonize almost every aquatic and terrestrial ecosystem (Lynn, 2008), including those pristine and highly impacted ones (Foissner and Berger, 1996), indicating adaptations to tolerate or even to degrade and metabolize environmental contaminant. Since many species are fast-growing and culturable under *in vitro* conditions and some of them are amenable to genetic modification approaches a great (yet underexplored) biotechnological potential is foreseen. In fact, six new tyrosinase homologs from four freshwater culturable ciliate species were identified and characterized using bioinformatics. Theoretical 3D models were generated, using comparative modeling, and the spectrum of action of these new tyrosinases were predicted against a set of ~1000 CECs structures through molecular docking (virtual screening) assays. Our data indicate that these tyrosinases of ciliates may target a wide variety of CECs, including antibiotics, licit and illicit drugs, pharmaceuticals, cosmetics, personal care products, cyanotoxins and pesticides, expanding the potential environmental applications of these versatile enzymes.

## 2. Materials and Methods

### 2.1. Genomic screenings and primary structure characterization of tyrosinase homologs

Draft genomes of 23 ciliate species were retrieved from GenBank (https://www.ncbi.nlm.nih.gov/genbank/) and screened for the presence of tyrosinase homologs. For each genome, open reading frames (orfs) (>300bp) and putative proteomes were inferred using *getorf* (Rice et al., 2000), applying the appropriate genetic codes, as suggested in analyses using *facil* (Dutilh et al., 2011). To identify tyrosinase homologs within these predicted proteomes two complementary approaches were used: pairwise sequence alignments, using *blastp* v2.6 (Camacho et al., 2009) against protein sequences from SwissProt (https://www.uniprot.org/), applying an e-value cutoff of 10^−5^ and query coverage filter of 80%; and profile searches, using *fuzzpro* (Rice et al., 2000) with conserved tyrosinase signatures from Prosite (Sigrist et al., 2012), and *Hmmer* v3 (Eddy, 2011) with Hidden Markov Models (HMMs) profiles of tyrosinases from PFAM (El-Gebali et al., 2019). Candidates were, subsequentially, characterized using *InterProScan* v5 (Jones et al., 2014), *Phobius* (Käll et al., 2004) and *SignalP* v5 (Almagro Armenteros et al., 2019) for detection of conserved domains, presence of transmembrane terminals and secretion signals, respectively. Moreover, molecular weight (MW) and isoelectric points (pI) were estimated using *compute pI/Mw* (https://web.expasy.org/compute_pi/).

### 2.2. Phylogenetic analysis

The dataset used to infer phylogenetic relationships among the newly identified tyrosinases and previously characterized ones were generated using all amino acid sequences available from SwissProt (https://www.uniprot.org/) (consulted in Feb 2020), excluding fragment of sequences, and using the tyrosinase from *Rhizobium meliloti* (SwissProt:P33180) as outgroup. The dataset was, then, aligned using *MAFFT* v7 (Rozewicki et al., 2019), with default parameters, resulting a matrix of 39 sequences and 1207 characters. To determine the best model of sequence evolution within this dataset, *SMS* (Lefort et al., 2017) was used, and a maximum likelihood phylogenetic tree was inferred using *phyml* v7 (Guindon et al., 2010), applying the WAG +G +I model of sequence evolution, and branch support values was defined through the nonparametric bootstrap method of 1,000 pseudo-replicates.

### 2.3. Three-dimensional (3D) structure prediction of tyrosinase homologs

Comparative modeling, using *Modeller* v9.19 (Webb and Sali, 2016) were performed to predict the theoretical 3D structures of tyrosinase homologs of ciliates. Best templates for each homolog sequence were identified using *HHpred* (Zimmermann et al., 2017), which is defined by high-quality pairwise amino acid sequence alignments between subjects and query proteins (with known 3D structures) deposited in the Protein Data Bank (PDB) (Berman et al., 2000). 100 models were generated for each homolog sequence and the one with the lowest DOPE (Discrete Optimized Protein Structure) score were selected for further analyses. Finally, the quality of the best theoretical models were evaluated using *MolProbity* (Chen et al., 2010). Visualization and manipulation of all 3D models were done using UCSF *Chimera* v1.14 (Pettersen et al., 2004).

### 2.4. *Molecular dynamics* (MD) simulations

To evaluate the stability of the generated 3D models of tyrosinases of ciliates, we performed MD simulations using *Gromacs* v5 (Abraham et al., 2015), applying OPLS-AA force field (Jorgensen et al., 1996). Models were centered into dodecahedral boxes with sizes of at least 2.0 nm, from the surface of the model to the side of the box. Systems were solvated using SPC/E water model (Mark and Nilsson, 2001) and neutralized with 0.15 mol/L sodium chloride. After a steepest *descent* minimization of 50,000 steps using position restraints, systems were equilibrated, sequentially, using NVT followed by NPT, considering temperature of 300 K and pressure of 1 bar, each ensemble for 100 ps. Then, models were released from their position restraints and the production steps ran for 50 ns using *leap-frog* as integrator. All-atom bond lengths were linked using *LINCS* (Hess et al., 1997). Electrostatic interactions and van der Waals were calculated by using particle-mesh Ewald (PME) method (Essmann et al., 1995) with cutoffs of 1.0 nm. For each atom, the list of neighbors was updated every 2 fs. The stability of the models during the run was accessed through root-mean-square deviation (RMSD) and radius gyrase plots, and secondary structures of the models were assigned by means of Dictionary of Protein Secondary Structures (DSSP), using the Gromacs tool, do_dssp.

### 2.5. Virtual screenings

Potential targets of the new tyrosinases of ciliates were predicted using a molecular docking (virtual screening) approach. For that, binding energies (-kcal/mol) between tyrosinase 3D models and a set of 76 CECs, including pharmaceuticals, illicit drugs, chemicals and natural toxins were estimated. Two know ligands of tyrosinases, L-tyrosine (CID6057) and odopaquinone (CID), and a well-studied tyrosinase isolated from the basidiomycete mushroom, *Agaricus bisporus* (PDB:2y9wA) were used to serve as a reference to determine likely and unlikely targets.

Before the assay, putative cofactors and pockets were predicted using *COACH* (Yang et al., 2013, 2012) and *DoGSiteScorer* (https://proteins.plus/), respectively. Then, correct protonation states of these new tyrosinases were inferred using *Protein Plus* (https://proteins.plus/). Three dimensional CECs structures were retrieved from PubChem (https://pubchem.ncbi.nlm.nih.gov/) and when unavailable, 2D structures (in SMILE format) were downloaded, instead, and converted to 3D using *Obabel* (O’Boyle et al., 2011). All ligands were desalted, polar hydrogens added, and clashes and torsions resolved by minimization (steepest descent), using *Obabel* (O’Boyle et al., 2011). Next, grids of 10 Å were placed into the active type-3 copper center of the *oxy*-tyrosinases using *ADT* (<u>A</u>uto <u>D</u>ock <u>T</u>ools) (Morris et al., 2009), and then, an in-house python script was used to run the virtual screenings using *AutoDock Vina* (Trott and Olson, 2010) and to compile the results from the highest binding energies values of the best poses for each tyrosinase 3D models and ligand structures.

To further evaluate the use of these new tyrosinases to target pesticides within the environment, ~1000 pesticide structures from PubChem were subjected to a new round of virtual screening assay, but now running the analysis within the online server DockThor (https://dockthor.lncc.br/v2/), with default settings and following the same framework for protein and ligand preparation as mentioned above.

## 3. Results and Discussion

### 3.1. Identification and primary structure characterization

The identification of tyrosinase homologs from the inferred proteome of 23 ciliate species was done using a combined approach based in pair-wise alignments against protein sequences from the manually curated protein database, Swiss-Prot; and profile-based searches using tyrosinase HMM-profiles from PFAM and tyrosinase signatures from PROSITE (as described in details in Method section). Using this framework, we identified six new tyrosinase homologs within the genomes of four freshwater stichotrichids, *Paraurostyla* sp. (PuTYR-1), *Stylonychia lemnae* (SlTYR-1 and SlTYR-2), *Uroleptopsis citrina* (UcTYR-1 and UcTYR-2) and *Urostyla* sp. (UTYR-1). These data are summarized in Table 1.

**Table 1.** Summarized data and general features of the identified tyrosinanes of ciliates.

| Putative tyrosinase | Species | NCBI accession number | Genomic position (nt) | BLASTP (SwissProt) |  |  | Length (aa) | Tyrosinase conserved domain (InterPro) | General features |  |  |  |
| --- | --- | --- | --- | --- | --- | --- | --- | --- | --- | --- | --- | --- |
| | | | | ϕHSP | Identity (%) | Host | | | Phobius | SignalP | Molecular weight (KDa) | Isoelectric point |
| PuTYR-1 | *Paraurostyla* sp. PUJRC_G6 | LASR02004360 | Contig 4360 (127 – 1713) | Tyrosinase (Q00234) | 39,69 | *Aspergillus oryzae* | 529 | IPR002227<br>IPR008922 | Non-cytoplasmatic | - | 59,87 | 5,05 |
| SITYR-1 | *Stylonychia lemnae* | CCKQ01010535 | Contig 0171 (2469 – 4043) | Tyrosinase (Q00234) | 39,21 | *Aspergillus oryzae* | 525 | IPR002227<br>IPR008922 | Non-cytoplasmatic | - | 59,66 | 5,08 |
| SITYR-2 | *Stylonychia lemnae* | CCKQ01013503 | Contig 0533 (1717 – 137) | Tyrosinase (Q00234) | 39,42 | *Aspergillus oryzae* | 527 | IPR002227<br>IPR008922 | Non-cytoplasmatic | - | 59,8 | 4,99 |
| UcTYR-1 | *Uroleptopsis citrina* | LXJT01005355 | Contig 3255 (1651 – 107) | Tyrosinase (Q00234) | 39,47 | *Aspergillus oryzae* | 515 | IPR002227<br>IPR008922 | Non-cytoplasmatic | - | 58,24 | 5,99 |
| UcTYR-2 | *Uroleptopsis citrina* | LXJT01021807 | Contig 3230 (116 – 1657) | Tyrosinase (Q00234) | 39,56 | *Aspergillus oryzae* | 514 | IPR002227<br>IPR008922 | Non-cytoplasmatic | - | 58,08 | 5,95 |
| UTYR-1 | *Urostyla* sp. PUJRC_G1 | LASQ02000317 | Contig 0317 (2405 – 72) | Tyrosinase (Q00234) | 38,84 | *Aspergillus oryzae* | 523 | IPR002227<br>IPR008922 | Non-cytoplasmatic | - | 60,5 | 6,33 |

$\phi$ HSP - High Scoring Pair

These tyrosinase homologs have a sequence identity of 39% with a tyrosinase characterized from the filamentous fungus *Aspergillus orizae* (SwissProt:Q00234) and carry conserved tyrosinase domains (InterProt domains IPR002227 and IPR008922) (Table 1). Their expected molecular weight (MW) and isoelectric point (pI) are within the range of typical tyrosinases described form literature (MW 14-75 KDa; pI 4.65 and 11.9) (Agarwal et al., 2016; Ba and Vinoth Kumar, 2017) and interestingly, all of them were predicted as extracellular enzymes, but no clear secretory signals could be identified (Table 1). This character was previously reported in some fungal tyrosinases (Mayer, 2006) and a possible route for their secretion would be through quaternary associations with “caddy-like” proteins caring their own signal peptides, such as those described in *Streptomyces* sp. (Matoba et al., 2018). The sequence similarity between tyrosinases of ciliates and fungi is further evidenced in our phylogenetic reconstruction (Figure 1), the ciliate`s homologs emerged in a single cluster together with the tyrosinase from *A. orizae* (SwissProt:Q00234) (Ciliophora/Fungi Clade I) that is a sister group of tyrosinases from higher fungi (Fungi Clade II), apart from tyrosinase clusters of *Streptomyces* (Bacteria Clade), insects (Metazoa Clade I), and mammals (Metazoa Clade II) (Figure 1).

**Figure 1.**
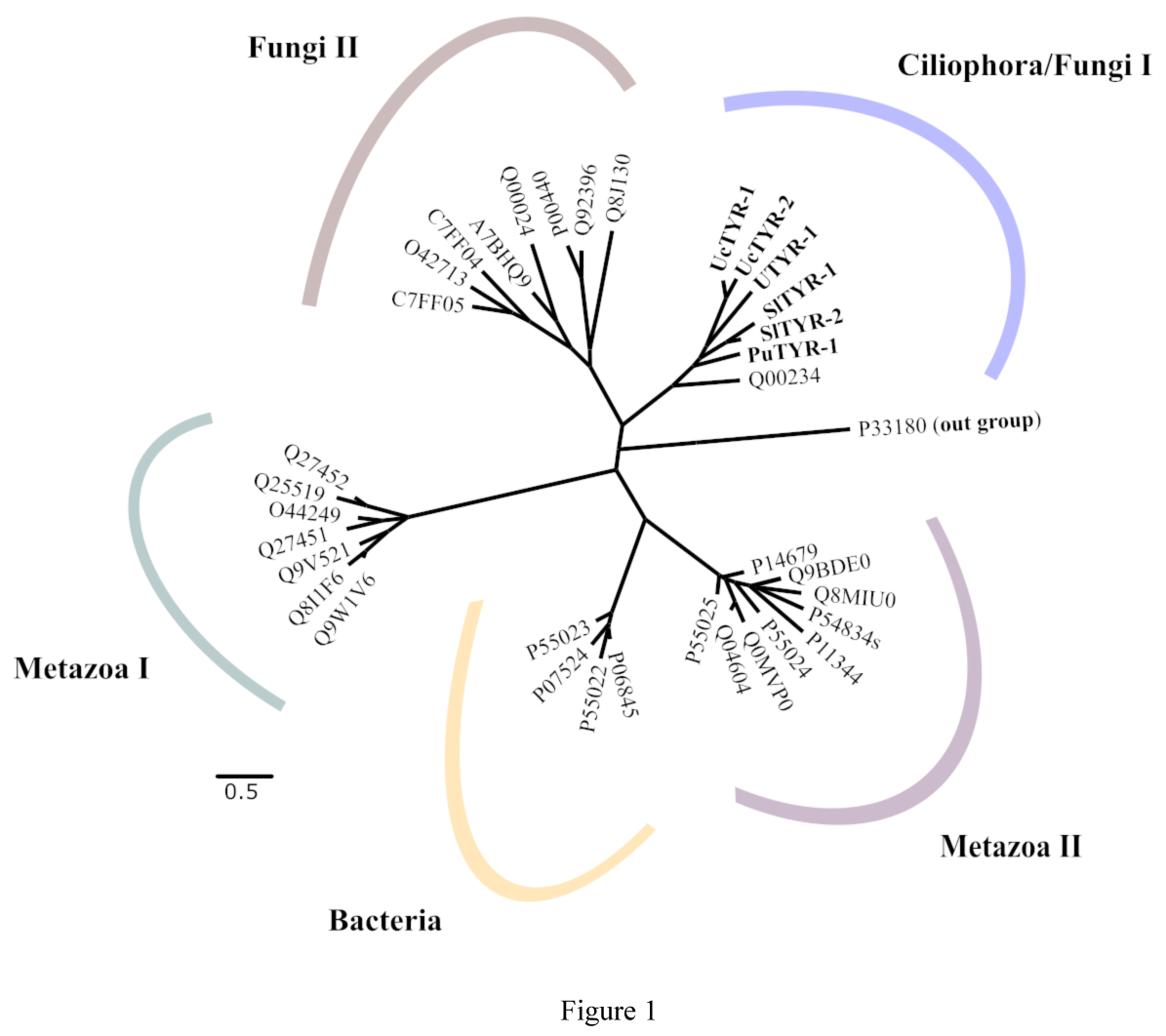
Phylogenetic reconstruction of tyrosinases based on amino acid sequences and inferred using maximum likelihood framework. Tyrosinases from ciliates are in bold. Barr represent the number of substitutions per 100 aa.

### 3.2. Tertiary structure prediction

We used the X-ray diffraction-defined structure of a tyrosinase isolated from *Burkholderia thailandensis* (PDB:5ZRD) as template to predicted, through comparative modeling, the theoretical 3D models of these tyrosinase homologs of ciliates (Table 2 and Figure 2). Their overall quality are rather satisfactory, >97% of amino acid residues are in allowed positions (Ramachandran plots) and low Molprobity index values, which reflects the crystallographic resolution at which each model is expected based in combined information derived from the clash score, percentage of unfavored residues within Ramachandran plot and bad side-chain rotamers (Table 2). These 3D models are pearl-shaped structures supported by a series of 8 α-helices secondary structures (Figure 2) with the characteristic active site, in central position, consisting of two copper ions (CuA and CuB), each one linked to three conserved histidine residues (Figure 3), within a pocket of mean volume of 411.43 Å^3^ and surface of 523.44 Å^2^ (Table 2). Five out six conserved histidine residues are rigids and located within α-helical structures, while the sixth is flexible and located within a loop region (Figure 3). We could also detect within the active site of these enzymes: a conserved cysteine residue, typical of type-3 eukaryotic tyrosinases (Gielens et al., 1997), which interacts with the flexible histidine residue, through covalent thioester bonding, to favor copper ions incorporation into the active site (Fujieda et al., 2013); a conserved asparagine residue, possibly involved in hydrogen-bonding interactions during monophenol deprotonation reactions (Goldfeder et al., 2014); and 2 bulk phenylalanine residues close to CuA, which may be associated with monophenolase activity hindering (Mauracher et al., 2014) (Figure 3).

**Table 2.** Structural analyses of ciliate’s tyrosinases.

| Putative tyrosinase | Template | ID. (%) | Cov. | Ramachandran plot |  |  | MolProbity score† | COACH |  | DoGSiteScorer |  | 5'UTR (aa) | C-Terminal Propeptide (aa) |
| --- | --- | --- | --- | --- | --- | --- | --- | --- | --- | --- | --- | --- | --- |
|  |  |  |  | Favorable positions (%) | Allowed position (%) | Outliers |  | Cof. | C-score | Pocket Volume (Å<sup>3</sup>) | Pocket Surface (Å<sup>2</sup>) |  |  |
| PuTYR-1 | 5ZRD C | 52,24 | 0,95 | 95,7 | 99 | 4 | 2,64 | Cu<sub>2</sub>O<sub>2</sub> | 0.24 | 172.93 | 250.73 | 1-4 | 405-529 |
| SITYR-1 | 5ZRD C | 48,4 | 0,96 | 96,4 | 99 | 4 | 0,79 | Cu<sub>2</sub>O<sub>2</sub> | 0.19 | 319.30 | 359.05 | 1-4 | 396-525 |
| SITYR-2 | 5ZRD C | 53,05 | 0,96 | 93,1 | 97,4 | 10 | 1 | Cu<sub>2</sub>O<sub>2</sub> | 0.27 | 202.42 | 270.30 | 1-4 | 398-527 |
| UcTYR-1 | 5ZRD C | 51,73 | 0,96 | 95,9 | 99,5 | 2 | 2,46 | Cu<sub>2</sub>O<sub>2</sub> | 0.26 | 262.85 | 389.62 | 1-4 | 396-515 |
| UcTYR-2 | 5ZRD C | 50,4 | 0,97 | 94,3 | 98,4 | 6 | 1,4 | Cu<sub>2</sub>O<sub>2</sub> | 0.23 | 854.21 | 1160.99 | 1-4 | 395-514 |
| UTYR-1 | 5ZRD C | 52,52 | 0,95 | 94,9 | 97,7 | 9 | 2,53 | Cu<sub>2</sub>O<sub>2</sub> | 0.23 | 656.90 | 709.97 | 1-4 | 399-523 |

†MolProbity score is a single score which combines the clashscore, rotamer, and Ramachandran evaluations. C-score is the confidence score of the COACH prediction. ID. - Identity. Cov. - Coverage. Cof. - cofactor

**Figure 2.**
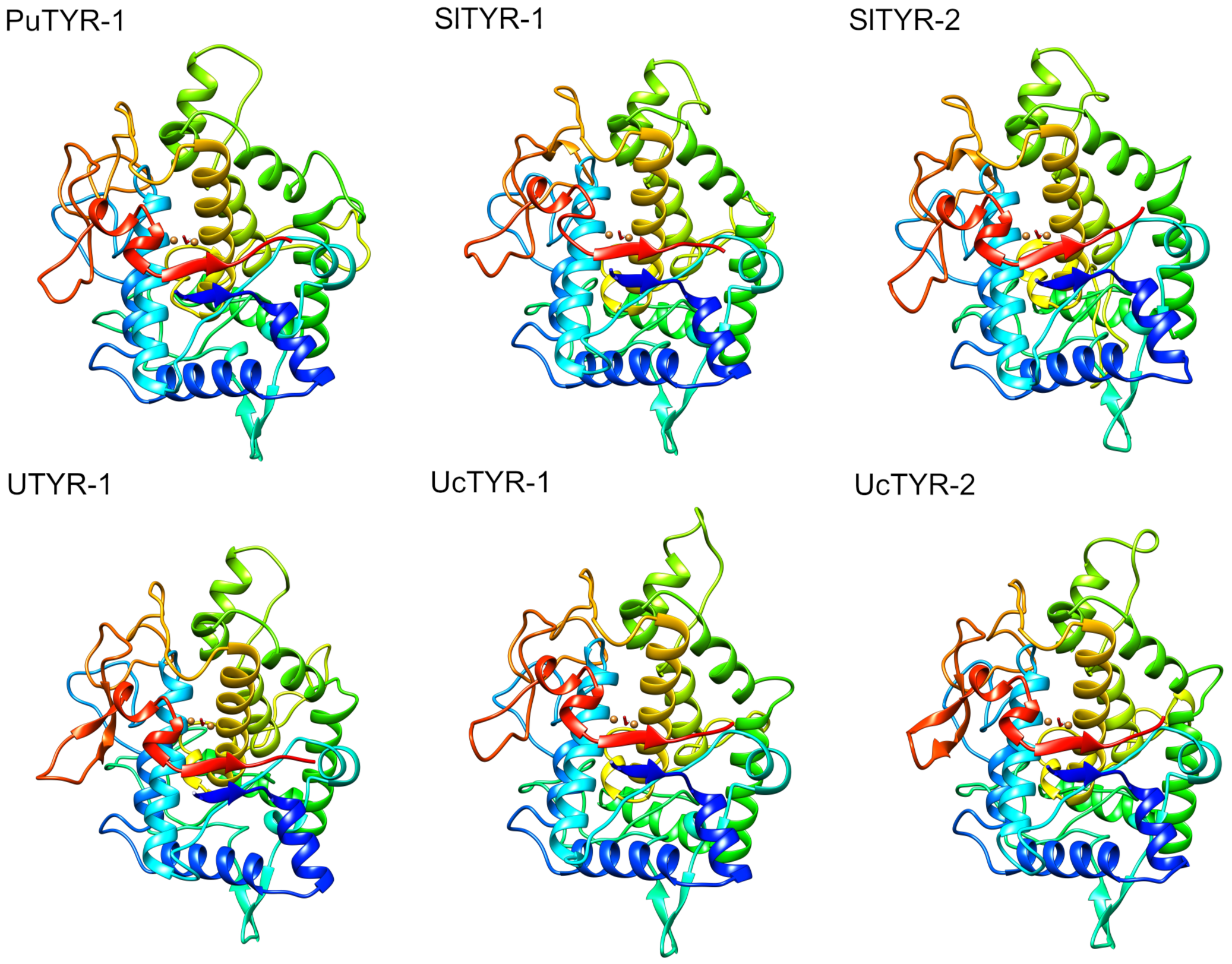
Three-dimensional structures of the six tyrosinases from ciliates (PuTYR-1, SlTYR-1, SlTYR-2, UcTYR-1, UcTYR-2 and UTYR-1). The two golden spheres represent copper atoms and the red rod, a dioxygen molecule.

**Figure 3.**
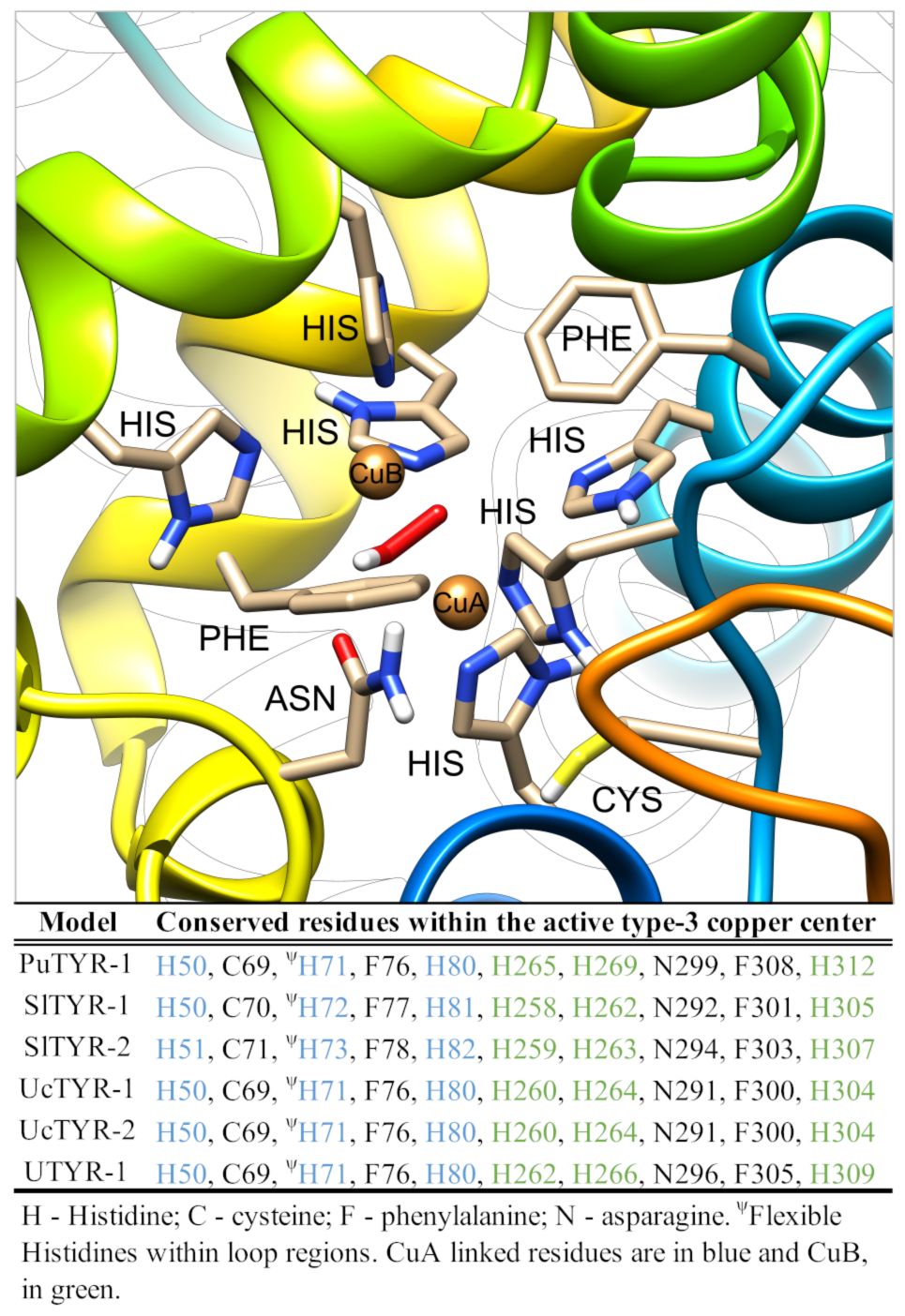
Representative active site of tyrosinases of ciliates based on the structure of PuTYR-1. Copper center formed by CuA and CuB (golden spheres), dioxygen (red rod), and conserved asparagine, histidine, phenylalanine, cysteine residues are presented.

These new tyrosinases seems to be synthesized in a latent form, which requires post-translational modifications to become active. They all present a C-terminal region of ~120 aa that is absent from well-characterized mature fungi tyrosinases, such as PDB:4OUA and PDB:2Y9W (Table 2), which, when present, covers the active site of the enzyme, possibly a strategy to control enzymatic activity. Moreover, these homologs present a short N-terminal region (~4 aa), with no significant alignment with any known protein from PDB, possibly representing a 5′ UTR. Both regions were. Both regions were trimmed off from the tyrosinases before modeling their 3D structures to avoid artifact problems in the subsequent docking analyses.

Before performing the molecular docking assay to predict the spectrum of action of these new tyrosinases against CECs, we subjected these theoretical models to molecular dynamics (MD) simulations of 50 ns to access their stability. Data were evaluated through root mean square deviation (RMSD) and radius of gyration plots (Figure 4), to reveal variations in the backbone and in the globularity of each one of these models, respectively, and by DSSP analyses (Suppl. Figure 1), to evidence fluctuations within their secondary structures during the MD simulation. Accordingly, the models showed, in general, low variations within backbone (Figure 4a), only slight changes into their globularity (Figure 4b), and their overall secondary structure organization remained considerably uniform (Suppl. Figure 1) along the MD run of 50 ns. The highest variations were observed for SlTYR-1, which varied 4,50 nm in the RMSD and 2,17 nm in the globularity. However, such variations could be considered within acceptable ranges for the analyses of potential spectrum of action over CECs.

**Figure 4.**
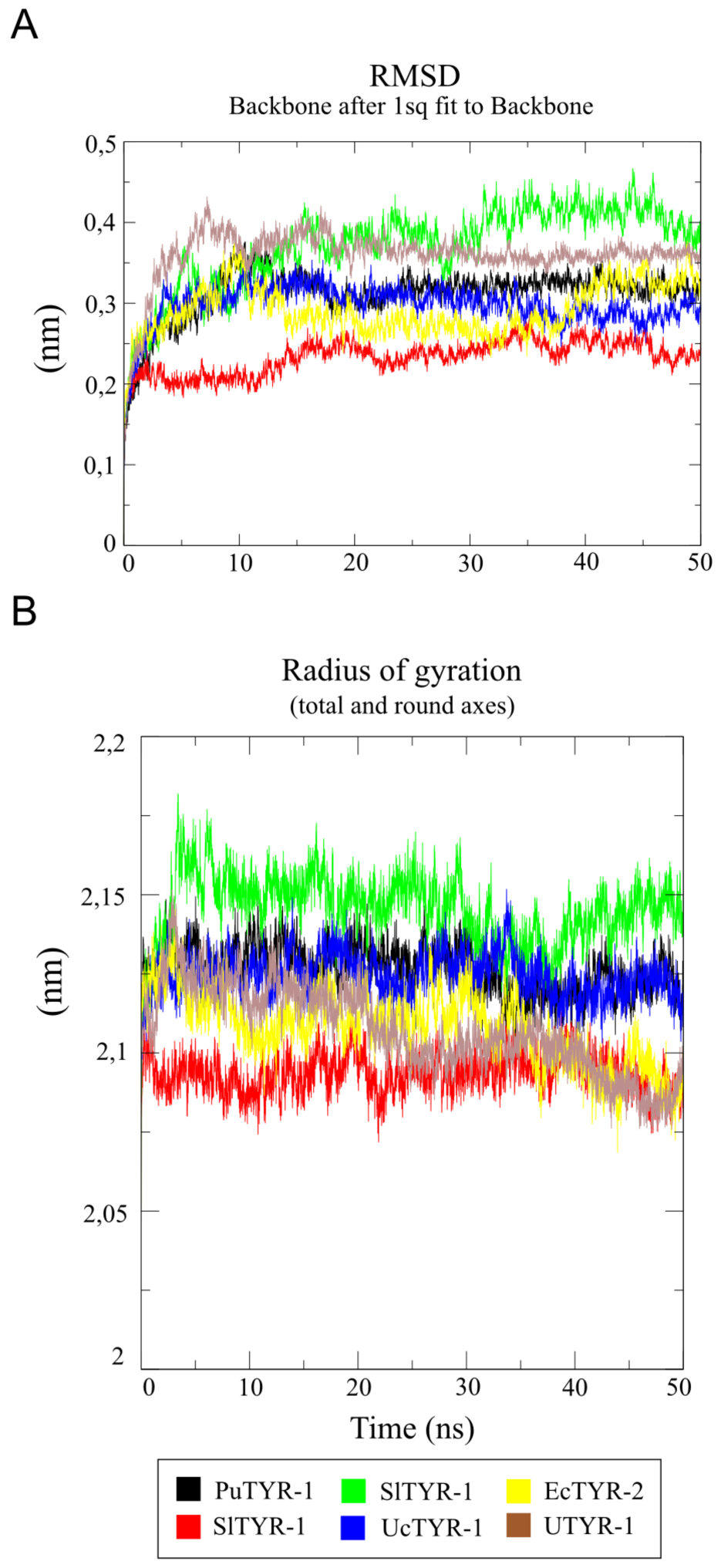
Molecular dynamic evaluation of PuTYR-1, SlTYR-1, SlTYR-2, UcTYR-1, UcTYR-2, and UTYR-1 theoretical structures. (A) a root mean square deviation (RMSD) plot, showing the mean backbone variation over the 50 ns of run; and (B) a radius of gyration plot, showing the mean variation within the globularity of each structure over the 50 ns of run.

### 3.3. Spectrum of action prediction

The spectrum of action of these new tyrosinase homologs of ciliates might be much broader than previously thought (Figure 5, Table 3). To evaluate potential targets for these enzymes, we first estimated binding energies between the generated 3D models and a set of 76 CECs structures, including antibiotics, industrial chemicals, illicit drugs, pesticides, pharmaceuticals, and natural toxins. Our rationale was that ligands with high binding energies, above the positive controls (bottom part of Figure 5), may represent positive interactions. As another second reference, we included the well-studied fungus tyrosinase PDB:2Y9WA (left side of Figure 5) in the analysis, and its general behavior was greatly similar to the tyrosinases from ciliates. As shown in Figure 5, binding energies vary considerably within tyrosinases and within ligands, ranging from −3,8 kcal/mol, between acephate and UcTYR-1, to −9,0 kcal/mol between microcystin and SlTYR-1, but in general, high binding energy values were obtained for most of the tested classes, especially for antibiotics, cyanotoxins, drugs, and hormones, where the highest values were −7.6 kcal/mol between UTYR-1 and the antibiotic ofloxacin (CID 4583); −9.0 kcal/mol between SlTYR-1 and the cyanotoxin Anatoxin-a (CID 3034748), −7.4 kcal/mol UTYR-1 and the pharmaceutical drugs Carbamazepine (CID 2554) and Diazepam (CID 3016); and −7.8 kcal/mol between UTYR-1 and the hormone estrone (CID 5870). On the other hand, the class pesticides, as whole, presented only low binding energy values, suggesting the group are unlikely targets for tyrosinases. However, given the environmental relevance of pesticides and to test whether the general low binding energies was specifically related with the selected pesticides (Figure 5), we decided to repeat the assay, now using ~1000 pesticide structures. The top 10 ligands with the highest binding energies are summarized in Table 3. In fact, high binding energies, way above the control, could be detected against a number of pesticides of different chemical groups, including widespread ones, such as the coumarininc rodenticide brodifacoum (CID 54680676); the organophosphorus herbicide falone (CID 7209), and the trifluoromethyl aminohydrazone insecticide hydramethylnon (CID 5281875) among many others (Table 3).

**Table 3.**
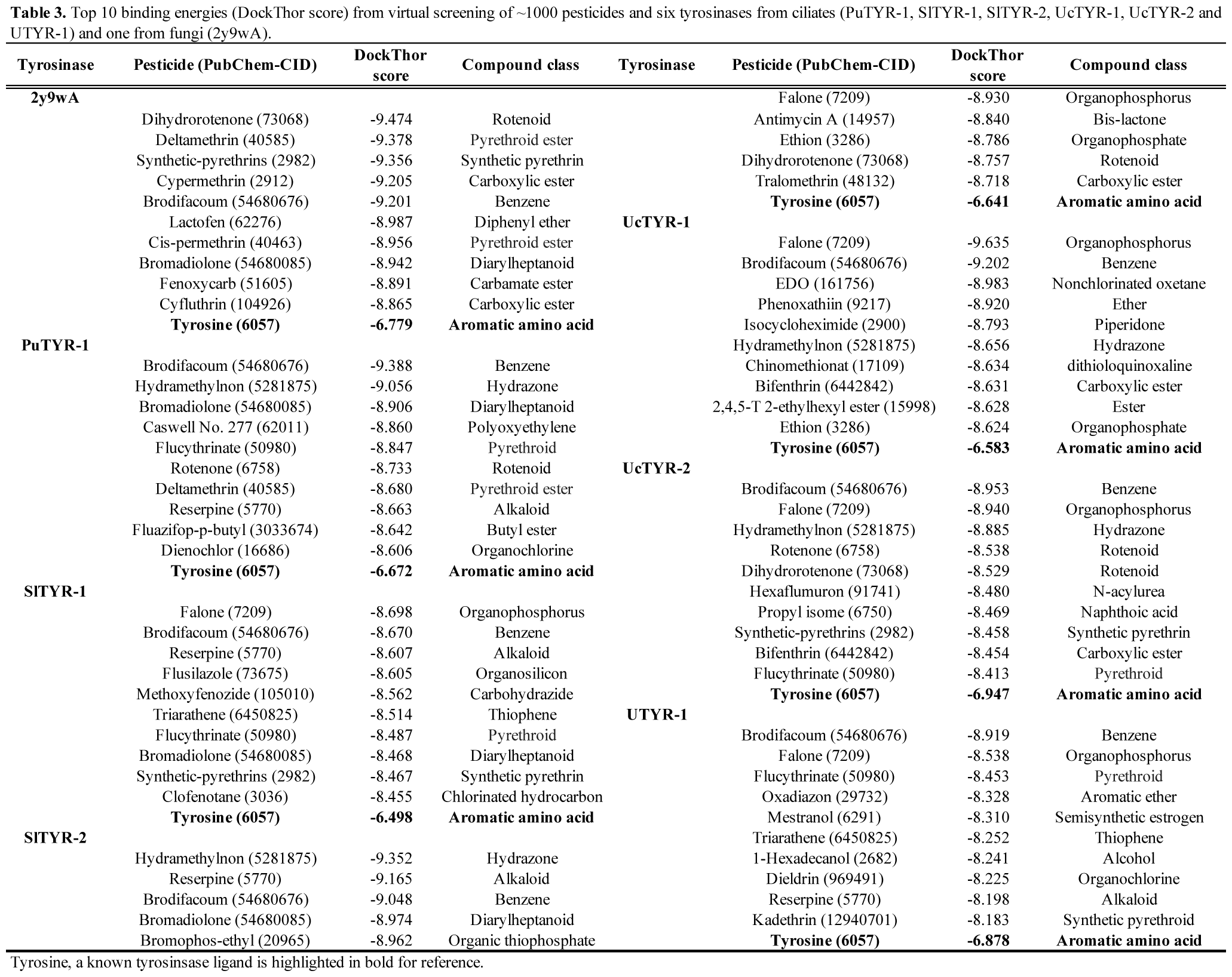
Top 10 binding energies (DockThor score) from virtual screening of ~1000 pesticides and six tyrosinases from ciliates (PuTYR-1, SlTYR-1, SlTYR-2, UcTYR-1, UcTYR-2 and UTYR-1) and one from fungi (2y9wA).

**Figure 5.**
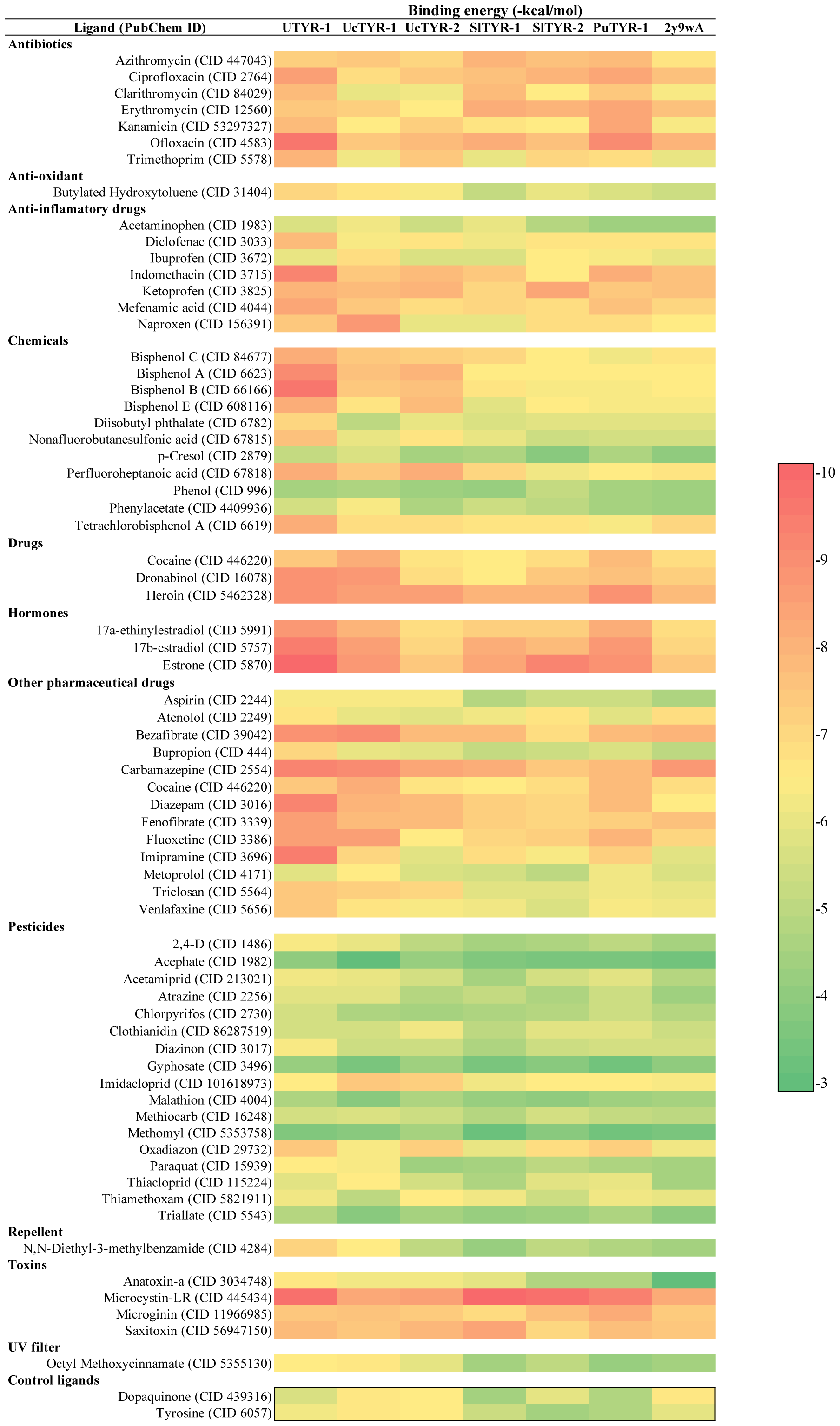
Binging energies (score) values from virtual screening of 76 CEC structures against six tyrosinases from ciliates (PuTYR-1, SlTYR-1, SlTYR-2, UcTYR-1, UcTYR-2 and UTYR-1) and one from fungi (2y9wA).

Interestingly, in both virtual screenings performed (Figure 5 and Table 3) binding energies varied considerably between tyrosinases and CECs, which could be explained by their high degree of polymorphism. In fact, pairwise sequence distances among PuTYR-1, SlTYR-1, SlTYR-2, UcTYR-1, UcTYR-2, and UTYR-1 are relatively high (0.46 in average), indicating numerous inter- and intraspecific polymorphisms (Suppl. Figure 2), which may have impact on their affinity to ligands. Contrasting binding energies were found, for example, for UTYR-1 (−7.5 kcal/mol) and UcTYR-2 (−5.5 kcal/mol) and Imipramine (CID 3696), an antidepressant drug that is associated with reduction in neuron numbers within the frontal lobe of fetal rats (Swerts et al., 2010); and for UTYR-1 (−7.6 kcal/mol) and PuTYR-1 (−5.8 kcal/mol) and Bisphenol B (CID 66166), a widely used plasticizer agent and know endocrine disruptor associated with different reproductive disorders in rats and fishes (Serra et al., 2019). Future studies aiming to map the influence of these polymorphisms in the binding energies against CECs will greatly contribute to the rational design of more efficient and ligand-specific tyrosinases.

Despite of our data indicate tyrosinases may have a broad spectrum of action against CECs, expanding the potential environmental applications of these versatile enzymes, all data generated require further experimental validation. Nevertheless, it is known from the literature that tyrosinases are effective for removal of the plasticizer agents and endocrine disruptors bisphenols A and B (Nicolucci et al., 2011), the methylphenol p-cresol (Bayramoglu et al., 2013) and Acetaminophen (Ba et al., 2014) from water matrices, which were also tested here; therefore, their binding energies would serve as references to guide future *in vitro* works toward the most likely targets for tyrosinases.

According to Ba et al. (Ba and Vinoth Kumar, 2017), despite the great biotechnological potential foreseen for tyrosinases in remediation and monitoring processes, high costs and low yield of purified enzymes are the main limitations for the widespread use of these enzymes. The six tyrosinases were identified four *in vitro* culturable and widespread freshwater ciliates, *Paraurostyla* sp. (PuTYR-1), *Stylonychia lemnae* (SlTYR-1 and SlTYR-2), *Uroleptopsis citrina* (UcTYR-1 and UcTYR-2), and *Urostyla* sp. (UTYR-1). Similarly, many ciliate species can be efficiently cultured under *in vitro* conditions, they have small sizes and fast generation times, and some of them (especially those from genera *Paramecium* and *Tetrahymena*) have a wide array of well-established molecular tools available. Moreover, numerous molecules of industrial relevance have been identified from a wide variety of ciliate species (Elguero et al., 2019), including tyrosinases (this work), antimicrobials (Petrelli et al., 2012), antitumoral (Carpi et al., 2018), and bioactive molecules (Anesi et al., 2016). With that, we predict a great potential in ciliates to reduce production costs and to serve as sources of novel molecules of industrial and pharmaceutical interest, but also of environmental relevance.

## 4. Conclusion

Tyrosinases (EC1.14.18.1) are bifunctional copper monooxygenases of prokaryotes and eukaryotes that have a great environmental biotechnological potential to monitor and transform a variety of phenolic and non-phenolic aromatic compounds. Here, six new tyrosinases were identified from the genome of four freshwater stichtrichid ciliates, *Paraurostyla* sp. (PuTYR-1), *Stylonychia lemnae* (SlTYR-1 and SlTYR-2), *Uroleptopsis citrina* (UcTYR-1 and UcTYR-2) and *Urostyla* sp. (UTYR-1) present conserved domains and the characteristic active type-3 copper center, of well-characterized tyrosinases, representing the first report of tyrosinases in protists. Based on the measured binding energies in the virtual screening assays, these new enzymes may bind, or even be active against antibiotics, anti-oxidants, anti-inflammatory drugs, industrial chemicals, illicit drugs, hormones, pharmaceutical drugs, insect repellents, cyanotoxins, UV filters, and pesticides from different classes, indicating that the spectrum of action and biotechnological potential of this family of enzymes against contaminants of emerging concern might be considerably broader than previously expected. Many ciliates, including these caring tyrosinase genes, have short generation times, can be easily *in vitro* cultivated, are amenable to genetic modifications and heterologous gene expression, and are known to produce a variety of secondary metabolites of biotechnological relevance. Therefore, we would like to stress the great biotechnological potential of ciliates as sources of novel molecules and to reduce costs of industrial-scale production of relevant enzymes. Although this is a pure bioinformatic work and requires proper biochemical data validation, it highlights the importance of a more intimate integration between bioinformatics and environmental biotechnology, which can greatly contribute to accelerate and reduce costs of new enzymes discovery/production steps for uses in remediation and monitoring processes of contaminants of emerging contaminants within natural environments.

## Acknowledgement

The authors acknowledge the “Coordenação de Aperfeiçoamento de Pessoal de Nível Superior” (CAPES) for the post-doctoral fellowship (PNPD) (88882.317976/2019-01) conferred to MS.

## Funding

This research did not receive any specific grant from funding agencies in the public, commercial, or not-for-profit sectors.

---

**Suppl. Figure 1.**
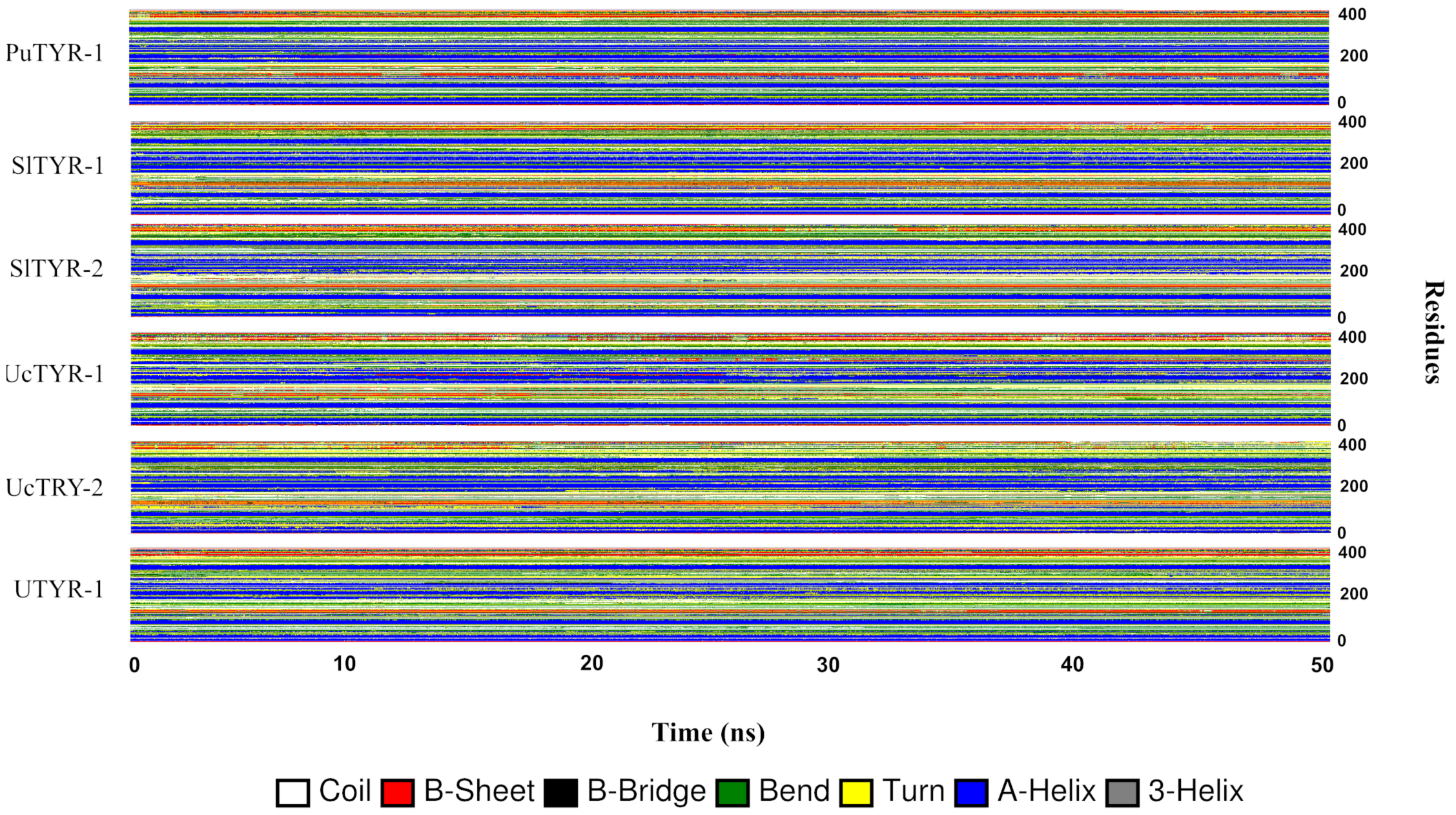
Secondary structure analyses (DSSP) of the models PuTYR-1, SlTYR-1, SlTYR-2, UcTYR-1, UcTYR-2 and UTYR-1 over the molecular dynamics run of 50 ns.

**Suppl. Figure 2.**
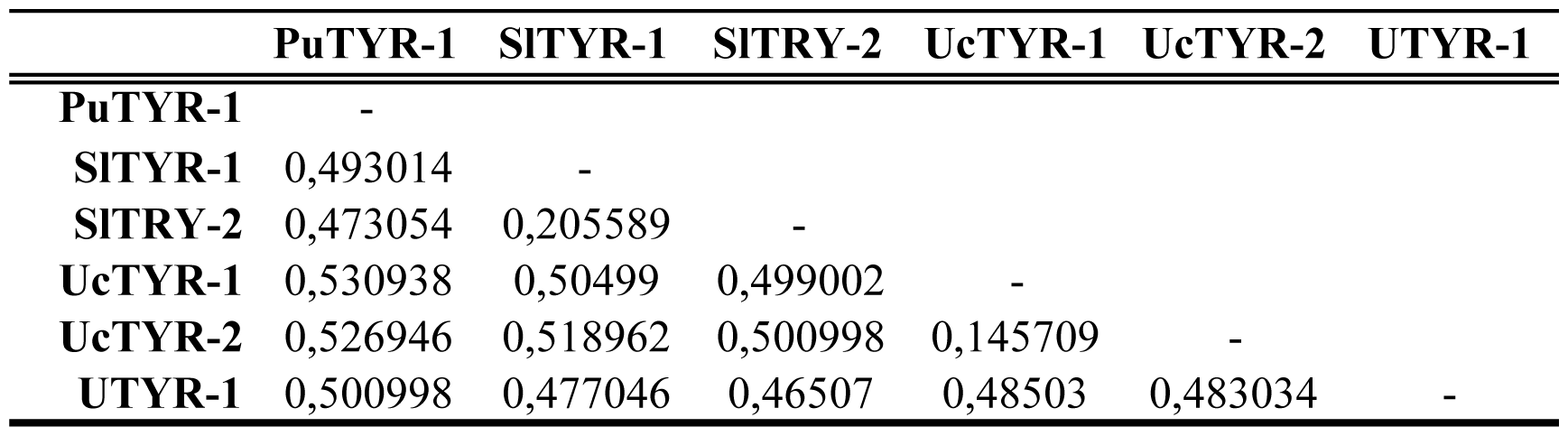
Pair-wise genetic distances between tyrosinases of ciliates.

## Notes

### Competing Interest Statement

The authors have declared no competing interest.

